# Immunogenicity of an Ad26-based SARS-CoV-2 Omicron Vaccine in Naïve Mice and SARS-CoV-2 Spike Pre-immune Hamsters

**DOI:** 10.1101/2022.03.04.482636

**Authors:** Maarten Swart, Adriaan de Wilde, Sonja Schmit-Tillemans, Johan Verspuij, Chenandly Daal, Ying Choi, Aditya Perkasa, Eleni Kourkouta, Issam Tahiri, Michel Mulders, Ana Izquierdo Gil, Leacky Muchene, Jarek Juraszek, Jort Vellinga, Jerome Custers, Rinke Bos, Hanneke Schuitemaker, Frank Wegmann, Ramon Roozendaal, Harmjan Kuipers, Roland Zahn

**Affiliations:** Janssen Vaccines & Prevention, Leiden, The Netherlands

**Author notes:** Corresponding author: Roland Zahn.

## Abstract

The severe acute respiratory syndrome coronavirus 2 (SARS-CoV-2) Omicron variant sparked concern due to its fast spread and the unprecedented number of mutations in the spike protein that enables it to partially evade spike-based COVID-19 vaccine-induced humoral immunity. In anticipation of a potential need for an Omicron spike-based vaccine, we generated an Ad26 vector encoding an Omicron (BA.1) spike protein (Ad26.COV2.S.529). Ad26.COV2.S.529 encodes for a prefusion stabilized spike protein, similar to the current COVID-19 vaccine Ad26.COV2.S encoding the Wuhan-Hu-1 spike protein. We verified that spike expression by Ad26.COV2.S.529 was comparable to Ad26.COV2.S. Immunogenicity of Ad26.COV2.S.529 was then evaluated in naïve mice and SARS-CoV-2 Wuhan-Hu-1 spike pre-immunized hamsters. In naïve mice, Ad26.COV2.S.529 elicited robust neutralizing antibodies against SARS-CoV-2 Omicron (BA.1) but not to SARS-CoV-2 Delta (B.1.617.2), while the opposite was observed for Ad26.COV2.S. In pre-immune hamsters, Ad26.COV2.S.529 vaccination resulted in robust increases in neutralizing antibody titers against both SARS-CoV-2 Omicron (BA.1) and Delta (B.1.617.2), while Ad26.COV2.S vaccination only increased neutralizing antibody titers against the Delta variant. Our data imply that Ad26.COV2.S.529 can both expand and boost a Wuhan-Hu-1 spike-primed humoral immune response to protect against distant SARS-CoV-2 variants.

## Introduction

In response to the SARS-CoV-2 (COVID-19) pandemic, multiple mostly spike-based vaccines were rapidly and successfully developed. These vaccines showed high efficacy against COVID-19 and have been deployed worldwide. Janssen developed the Ad26.COV2.S vaccine, which is a replication-incompetent human adenovirus type 26 (Ad26) vector^1^ encoding a stabilized pre-fusion SARS-CoV-2 spike protein based on the Wuhan-Hu-1 isolate^2^. A phase 3 trial demonstrated that Ad26.COV2.S was 74.6% efficacious at preventing severe-critical COVID-19^3^. The Ad26.COV2.S COVID-19 vaccine was granted emergency use authorization in the US and (conditional) marketing authorization in the European Union; in more than 50 other countries.

Several rapidly spreading SARS-CoV-2 variants have evolved since the initial introduction of the virus into humans. The Beta (B.1.351) and Delta (B.1.617.2) variants of concern (VOC) were initially thought to potentially evade vaccine elicited immunity. However, COVID-19 vaccine efficacy was largely maintained^3–8^. The Delta VOC obtained virtually worldwide dominance until it was replaced by the Omicron variant, BA.1 (initially named B.1.1.529). After its first reporting in November 2021, BA.1 was instantly declared a VOC due to its rapid spread and the unparalleled number of mutations in the spike protein. BA.1 carries 15 mutations in its receptor binding domain (RBD), which is an immunodominant target for neutralizing antibodies^9^, raising the possibility of reduced effectiveness of both vaccines and therapeutic monoclonal antibodies targeting this region^10–13^. Neutralizing antibody titers against SARS-CoV-2 Omicron BA.1 are indeed strongly reduced in vaccinees after the primary vaccination series compared to neutralizing antibody titers against the ancestral Wuhan-Hu-1 strain^14,15^. Importantly, a real-world evidence study showed that a homologous boost with Ad26.COV2.S administered 6-9 months after primary vaccination provided more than 80% protection against hospitalization during the Omicron wave in South Africa^16^. Vaccine protection against severe disease despite low Omicron neutralizing antibody titers may be maintained by conserved cellular immunity^17–19^ and non-neutralizing antibodies with Fc-functions^20^ across SARS-CoV-2 variants.

An Omicron (BA.1)-based vaccine candidate using the Ad26 vaccine platform (Ad26.COV2.S.529) was generated to assess whether the immunogenicity against the Omicron variant could be further improved. Here, we show that Ad26.COV2.S.529 induces robust Omicron neutralizing antibody titers in naïve mice and in hamsters with pre-existing SARS-CoV-2 spike protein immunity.

## Materials and methods

### Vaccines and challenge stocks

Replication-incompetent E1/E3-deleted adenovirus serotype 26 (Ad26) vector-based vaccines were generated using the AdVac system as described previously^2^. Ad26NCOV006 and Ad26.COV2.S encode a SARS-COV-2 spike protein sequence based on the SARS-CoV-2 Wuhan-Hu-1 spike protein (GenBank accession number MN908947), while Ad26.COV2.S.529 encodes a SARS-COV-2 spike protein sequence based on the SARS-CoV-2 Omicron BA.1 spike protein (GISAID accession number EPI_ISL_6913991). Spike proteins encoded by Ad26.COV2.S and Ad26.COV2.S.529 were stabilized in the prefusion conformation by the proline substitutions K986P and V987P and substitutions R682S and R685G, which abolish the furin cleavage site. The negative control vector Ad26.Empty, which did not contain a transgene, was used as a control.

### Cell-based ELISA

A549 cells were seeded at 2.9 × 10^4^ cells/well in Dulbecco’s Modified Eagle Medium (DMEM) with 10% heat-inactivated fetal bovine serum (FBS) in a flat-bottomed 96-well microtiter plate (Corning). The plate was incubated overnight at 37 °C in 10% CO^2^. After 24 h, cells were transduced with either Ad26.COV2.S or Ad26.COV2.S.529 at a dose of 2000 infectious units [IU]/cell and the cells were incubated for 48 h at 37 °C in 10% CO^2^. Two days post transduction, cells were washed four times with phosphate buffered saline (PBS) and subsequently fixed with 4% formaldehyde in PBS. After a 20-minute incubation at room temperature (RT), cells were washed four times with 0.05% Tween-20 in PBS. Cells were permeabilized by incubation with 1% Elugent (Merck) for 15-20 minutes, after which they were washed four times with 0.05% Tween-20 in PBS. Casein blocking buffer (Thermo Scientific) was then added and the cells were incubated for 60 to 90 minutes at 37 °C in 10% CO^2^. Cells were washed four times with 0.05% Tween-20 in PBS, after which two-fold diluted CV3-25 antibody or ACE2-Fc fusion protein (0.007 to 15 µg/ml) was added per well. CV3-25 was produced at ImmunoPrecise according to Jennewein et al.^21^ and ACE2-Fc was prepared according to Liu et al^22^. After 30 to 60 minutes of incubation, cells were washed four times with 0.05% Tween-20 in PBS. Next, cells were incubated with mouse HRP-conjugated anti-human IgG Fc (Jackson Immunoresearch, 1:8000) for 40 min at RT. The cells were washed four times with 0.05% Tween-20 in PBS. 3,3′,5,5′-Tetramethylbenzidine (TMB) Liquid Substrate System for ELISA (Sigma) was added and after 20 minutes the reaction was terminated by adding Stop Reagent for TMB Substrate (Sigma). Signal was measured at 450 nm (signal) and at 630nm (for background subtraction) using a Biotek Synergy Neo reader. Data were analyzed using GraphPad Prism 9, using the “Specific binding with Hill slope” module to calculate the antibody binding affinity.

### Animal studies

Animal experiments were approved by the Central Authority for Scientific Procedures on Animals (Centrale Commissie Dierproeven) and conducted in accordance with the European guidelines (EU directive on animal testing 86/609/EEC) and local Dutch legislation on animal experiments.

Female BALB/c mice aged 8-10 weeks at the start of study were purchased from Charles River Laboratories (Germany). Mice were immunized intramuscularly with 100 μl (50 μl per hindleg) vaccine under general anesthesia with isoflurane. Blood samples were collected via submandibular bleeding.

Female Syrian golden hamsters (*Mesocricetus auratus*), strain RjHan:aura, aged 9-11 weeks at the start of the study, were purchased from Janvier Labs (France). Hamsters were immunized intramuscularly with 100 μl (50 μl per hindleg) vaccine under general anesthesia with isoflurane. Blood samples were collected via the retro-orbital route under isoflurane anesthesia. Blood from all animal experiments was processed for serum isolation.

### Recombinant lentivirus-based pseudotyped virus neutralization assay (psVNA)

Neutralizing antibody titers were measured against several SARS-CoV-2 spike variants by a pseudotyped virus neutralization assay (psVNA). Human Immunodeficiency Virus (HIV)-based lentiviruses, pseudotyped with SARS-CoV-2 spike protein (based on Wuhan-Hu-1; GenBank accession number MN908947) were generated as described previously^23,24^. Substitutions and deletions in the spike protein open reading frame for the variant B.1.617.2^24^ and BA.1 (GISAID accession number EPI_ISL_6913991) were introduced using standard molecular biology techniques and confirmed by sequencing.

Assays were performed on HEK293T target cells stably expressing the human angiotensin-converting enzyme 2 (ACE2) and human transmembrane serine protease 2 (TMPRSS2) genes (VectorBuilder, Cat. CL0015). The cells were seeded in white half-area 96-well tissue culture plates (Perkin Elmer) at a density of 1.5 × 10^4^ cells/well.

Two-fold serial dilutions were prepared from heat-inactivated serum samples in phenol red free Dulbecco′s Modified Eagle′s Medium (DMEM) supplemented with 1% FBS and 1% PenStrep. The serially diluted serum samples were incubated at room temperature with an equal volume of pseudovirus particles with titers of approximately 1 × 10^5^ Relative Luminescence Units (RLU) luciferase activity. After one hour incubation, the serum-particle mixture was inoculated onto HEK293T.ACE2.TMPRSS2 cells. Luciferase activity was measured 40 h after transduction by adding an equal volume of NeoLite substrate (Perkin Elmer) to the wells according to the manufacturer’s protocol, followed by read out of RLU on the EnSight Multimode Plate Reader (Perkin Elmer). SARS-CoV-2 neutralizing titers were calculated using a four-parameter curve fit as the sample dilution at which a 50% reduction (N50) of luciferase readout was observed compared with luciferase readout in the absence of serum (High Control). The starting serum sample dilution of 20 was fixed as the lower limit of detection (LLOD).

### Statistical analysis

Statistical comparisons were performed in SAS 9.4 using a paired-sample t-test from an ANOVA. If titers were censored at LLOD, then a Tobit z-test from a Tobit ANOVA was used instead. If a vaccine-dose had more than 50% censored measurements, the non-parametric Mann-Whitney U-test was used instead. No adjustments for multiple comparisons were done.

## Results

### In-vitro characterization of spike expression by Ad26.COV2.S.529

Ad26.COV2.S.529 vector spike expression and antigenicity were characterized *in vitro* and compared to Ad26.COV2.S. Spike protein expression was evaluated after transduction of A549 cells using a quantitative cell-based ELISA with CV3-25 and ACE2-Fc. CV3-25 is an antibody that binds to the stem region of the SARS-CoV-2 spike S2^21^, which is conserved between the Wuhan-Hu-1 and BA.1 spike protein and ACE2-binding affinity is reported to be similar between Wuhan-Hu-1 Spike and Omicron spike^25,26^. Here, we show that CV3-25-binding to the spike protein expressed after transduction of A549 cells with Ad26.COV2.S or Ad26.COV2.S.529 was comparable (**Figure 1A**). Similarly, ACE2-Fc fusion protein binding to both spike proteins was comparable (**Figure 1B**).

**Figure 1.**
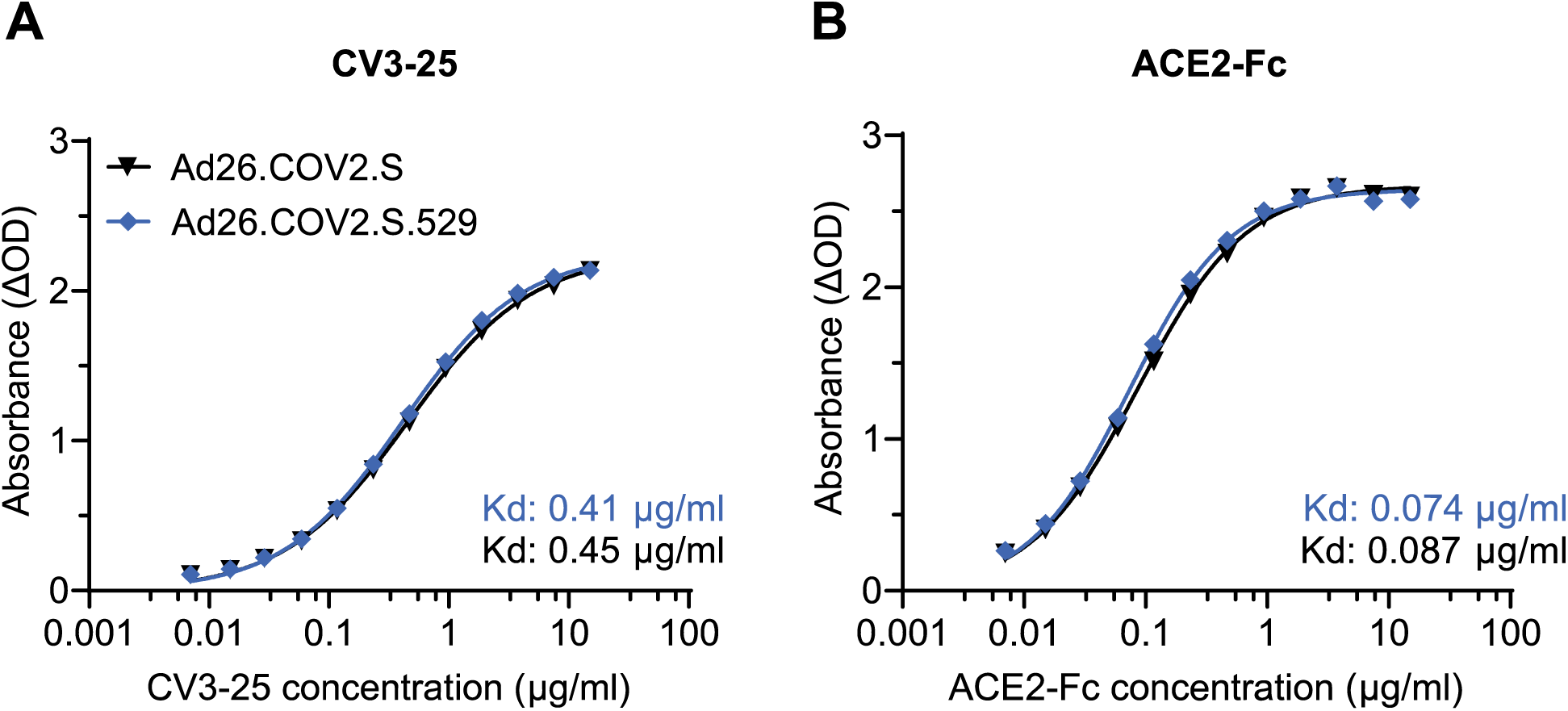
Comparison of CV3-25 and ACE2-Fc antibody binding to spike after expression by Ad26.COV2.S and Ad26.COV2.S.529. **A**. CV3-25 or **B**. ACE2-Fc fusion protein binding to the spike protein in A549 cells after transduction with 2000 infectious units/cell Ad26.COV2.S.529 (blue line) or Ad26.COV2.S (black line). Antibody binding affinity (Kd) values are shown, each point represents the mean value of a duplicate measurement.

### Ad26.COV2.S.529 induces Omicron neutralizing antibodies in both naïve mice and hamsters with pre-existing immunity

Naïve mice were immunized with 10^8^, 10^9^, 10^10^ viral particles (vp) of Ad26.COV2.S, Ad26.COV2.S.529 or 10^10^ vp Ad26.Empty mock control vector. SARS-CoV-2 spike neutralization titers were evaluated using a psVNA assay in sera collected 4 weeks after immunization. We focused our analysis on the two most recent and prevalent VOCs worldwide: Delta (B.1.617.2) and Omicron (BA.1). A single immunization with Ad26.COV2.S.529 induced dose-dependent Omicron spike neutralizing antibodies that were significantly higher than after vaccination with Ad26.COV2.S at all doses tested (**Figure 2A**). In contrast, Omicron spike neutralizing titers 4 weeks after vaccination with Ad26.COV2.S were comparable to animals vaccinated with Ad26.Empty. While Ad26.COV2.S induced robust dose-dependent Delta spike neutralizing antibody titers, Delta titers in animals vaccinated with 10^8^ and 10^9^ vp Ad26.COV2.S.529 were in the same range as after vaccination with Ad26.Empty (**Figure 2A**). Only the highest dose of Ad26.COV2.S.529 tested (10^10^ vp) induced statistically significantly Delta spike neutralization titers compared with Ad26.Empty, albeit at lower levels compared to 10^10^ vp Ad26.COV2.S.

**Figure 2.**
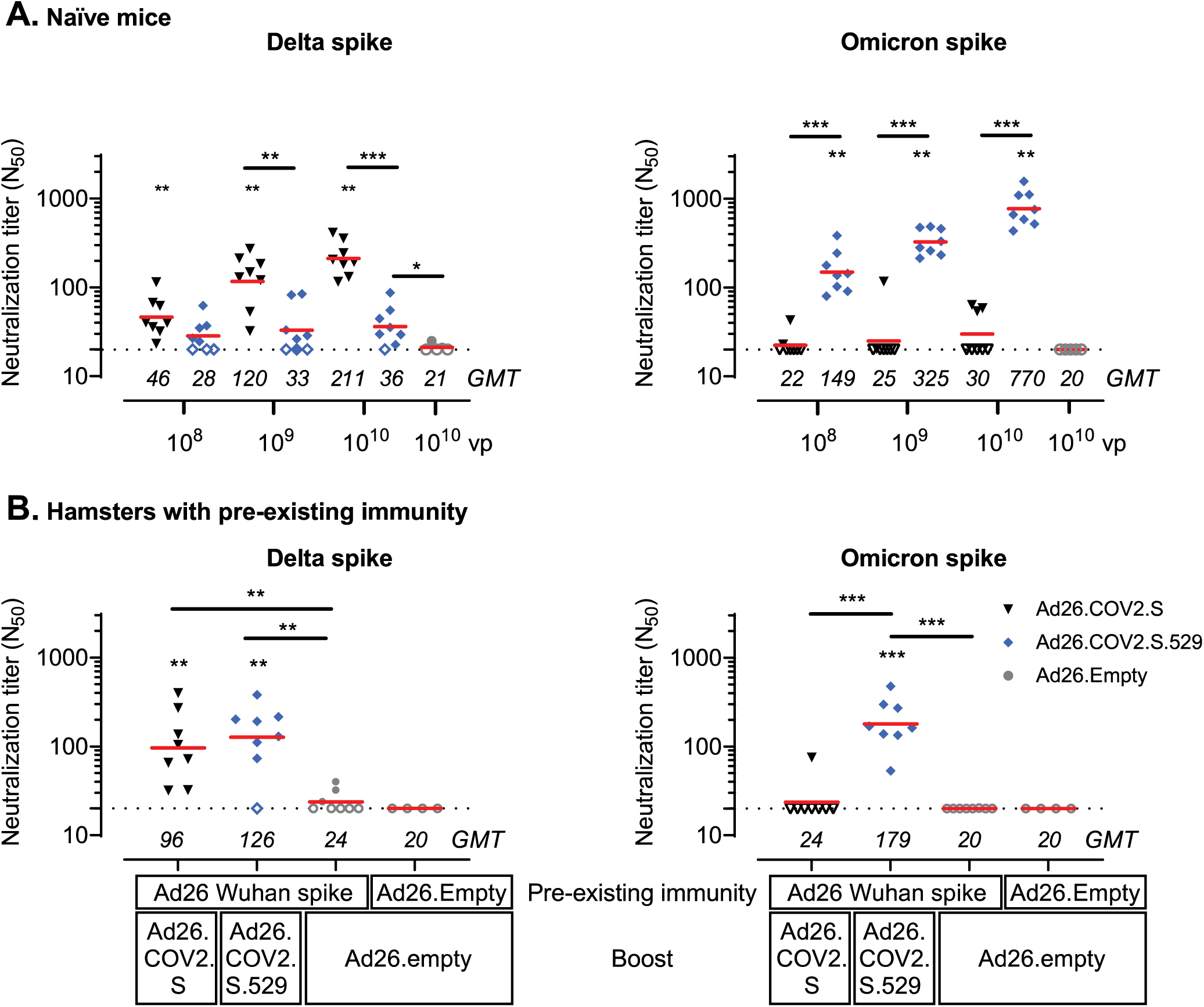
SARS-CoV-2 Delta and Omicron spike neutralizing antibodies induced by Ad26.COV2.S.529. **A**. Mice were immunized with 108, 109 or 1010 viral particles (vp) Ad26.COV2.S or Ad26.COV2.S.529 (both n=8); 1010 vp Ad26.Empty (n=5). Serum was collected 4 weeks after immunization to measure neutralizing titers against pseudotyped viruses expressing SARS-CoV-2 B.1.617.2 (Delta) or BA.1 (Omicron) spike in a pseudotyped virus neutralization assay (psVNA). **B**. Hamsters with pre-existing immunity were generated by vaccination with 107 viral particles (vp) of Ad26NCOV006 (Ad26 vector encoding Wuhan-Hu-1 spike) or Ad26.Empty. 6 weeks later hamsters were immunized with 1010 Ad26.COV2.S, Ad26.COV2.S.529 (both n=8) or Ad26.Empty (n=4). Serum was collected 4 weeks after this last immunization for psVNA analysis. Neutralizing antibody titers are expressed as the dilution giving a 50% reduction (N50) in the normalized luciferase readout (normalization relative to control wells without any serum added). Horizontal red bars and values per immunization group represent geometric mean (GMT) titers. Dashed horizontal lines represent the lower limit of detection (LLOD) of a 1:20 dilution, the lowest dilution measured in the assay. Open symbols indicate the response is at or below the LLOD. Pairwise comparisons were performed by a t-test, Tobit Z-test, or Mann-Whitney test. Significance is shown compared to the Ad26.Empty-immunized group unless depicted otherwise. *, P<0.05; **, P<0.01; ***, P<0.001.

As an increasing part of the population acquired pre-existing immunity either by infection or vaccination, we also evaluated the immunogenicity of Ad26.COV2.S.529 in hamsters with pre-existing immunity to a Wuhan-Hu-1 SARS-CoV-2 spike protein. Hamsters were first immunized with 10^7^ vp Ad26NCOV006, which encodes the non-stabilized ancestral SARS-CoV-2 Wuhan-Hu-1 spike protein. The mock control group was immunized with 10^7^ vp of Ad26.empty. Six weeks later, the hamsters received a vaccination with 10^10^ vp of Ad26.COV2.S, Ad26.COV2.S.529 or Ad26.Empty. SARS-CoV-2 Delta and Omicron spike neutralization titers were evaluated using a psVNA assay in sera collected 4 weeks after vaccination. Vaccination of pre-immune hamsters with Ad26.COV2.S and Ad26.COV2.S.529 resulted in comparable Delta spike neutralizing antibody titers that were significantly higher than in control animals (**Figure 2B**). While Omicron spike neutralizing antibody titers were undetectable in 7 out of 8 Ad26.COV2.S-vaccinated hamsters, vaccination with Ad26.COV2.S.529 induced robust Omicron spike neutralizing antibodies.

## Discussion

The SARS-CoV-2 Omicron variant carries an unprecedented number of mutations in the spike protein and a unique combination of mutations in the RBD that coincides with higher transmissibility and a substantial reduction in sensitivity to neutralizing antibodies elicited by natural SARS-CoV-2 infection or vaccination^13^. While a booster dose with current vaccines based on ancestral Wuhan-Hu-1 SARS-CoV-2 spike provides robust protection against Omicron infection-related severe disease^16,27^, protection from infection is lower and wanes faster, at least after an mRNA vaccine booster^28^. As a first step to evaluate if we can improve immunogenicity and efficacy against COVID-19 caused by the Omicron VOC, we generated an Ad26 vaccine candidate expressing a stabilized Omicron-based spike protein (Ad26.COV2.S.529) and evaluated its immunogenicity in comparison to Ad26.COV2.S in naïve mice and in a pre-immune hamster model.

We did not detect Omicron-neutralizing antibodies 4 weeks after immunization with Ad26.COV2.S in naïve mice and in hamsters with pre-existing immunity to ancestral Wuhan-Hu-1 spike. Although neutralizing antibodies against SARS-CoV-2 correlate well with vaccine-mediated protection in humans, cross-reacting cellular responses and non-neutralizing antibodies that exert Fc-mediated effector functions are likely contributing to protection as well. These immune functions seem largely conserved against Omicron^17–20^ and may have contributed to the effectiveness of a homologous booster with Ad26.COV2.S against hospitalization during the Omicron wave and to the effectiveness in macaques against a high-dose challenge with SARS-CoV-2 Omicron^16,29^.

Ad26.COV2.S.529, but not Ad26.COV2.S, induced robust Omicron (BA.1) spike neutralizing titers in naïve mice, in line with studies with Omicron-based RNA vaccines in naïve animals^30–34^. These differences in Omicron neutralizing titers are not attributed to differences in spike expression as we have shown that the expression of spike *in vitro* is comparable after transduction with Ad26.COV2.S and Ad26.COV2.S.529. In addition, Ad26.COV2.S.529 elicited robust Omicron (BA.1) spike neutralizing titers in pre-immune hamsters, while Ad26.COV2.S did not. The capacity of Ad26.COV2.S.529 to elicit antibodies with the ability to neutralize Omicron in hamsters that were pre-exposed to the spike protein of the SARS-CoV-2 Wuhan-Hu-1 strain indicates either boosting of low level pre-existing common neutralizing epitope specific memory B cells or de-novo induction of B cells that produce antibodies that can neutralize Omicron.

Our data seem to contrast data from an Omicron-based mRNA vaccine that provided similar neutralizing antibody levels against Omicron as compared to the original Wuhan-Hu-1-based mRNA vaccine in nonhuman primates (NHP) that were previously immunized with 2 doses of the original mRNA vaccine^35^. The Omicron-based mRNA vaccine did however elicit higher Omicron-neutralizing titers in pre-immune mice as compared to the original mRNA vaccine^31^. Nevertheless, we have previously shown in pre-immune NHPs, that a heterologous booster vaccination with an Ad26 vector-based vaccine candidate expressing a stabilized Beta-based spike protein (Ad26.COV2.S.351) >7 months after Ad26.COV2.S vaccination elicited 2.9-fold higher Beta psVNA titers than homologous vaccination with the Ad26.COV2.S^36^. It is currently unclear if these differences are related to the peak magnitude of immune responses elicited by previous immunizations, the number of pre-immunizations with the original vaccine prior to vaccination with the Omicron-based vaccine, the vaccine platform or the experimental animal models used.

Ad26.COV2.S.529 vaccination resulted in similar Delta spike neutralization titers as Ad26.COV2.S in pre-immune hamsters, but Delta neutralizing titers were undetectable or very low in Ad26.COV2.S.529 vaccinated naïve mice. The latter is consistent with data from naïve animals vaccinated with Omicron-specific mRNA vaccines^30–33^ and also in Omicron-infected hamsters^37^ where Delta spike neutralizing antibodies were undetectable. This suggests that an immunization with a heterologous spike protein as encoded by Ad26.COV2.S.529 induces a more broadly neutralizing antibody response in animals with pre-existing immunity to ancestral Wuhan-Hu-1 spike than a homologous spike protein immunization. Ongoing studies in non-human primates and potential future studies in humans may confirm these findings.

## Acknowledgements

We thank Mandy Jongeneelen, Remko van der Vlugt and Theo Schouten for providing the psVNA particles.

## References

1. Abbink, P. et al. Comparative seroprevalence and immunogenicity of six rare serotype recombinant adenovirus vaccine vectors from subgroups B and D. J. Virol. 81, 4654–4663 (2007).

2. Bos, R. et al. Ad26 vector-based COVID-19 vaccine encoding a prefusion-stabilized SARS-CoV-2 Spike immunogen induces potent humoral and cellular immune responses. NPJ Vaccines 5, 91 (2020).

3. Sadoff, J. et al. Final Analysis of Efficacy and Safety of Single-Dose Ad26.COV2.S. N. Engl. J. Med. 386, 847–860 (2022).

4. Polinski, J. M. et al. Effectiveness of the Single-Dose Ad26.COV2.S COVID Vaccine. MedRxiv (2021) doi:10.1101/2021.09.10.21263385.

5. Gier, B. de et al. COVID-19 vaccine effectiveness against hospitalizations and ICU admissions in the Netherlands, April-August 2021. MedRxiv (2021) doi:10.1101/2021.09.15.21263613.

6. Cohn, B. A., Cirillo, P. M., Murphy, C. C., Krigbaum, N. Y. & Wallace, A. W. SARS-CoV-2 vaccine protection and deaths among US veterans during 2021. Science (2021) doi:10.1126/science.abm0620.

7. Lopez Bernal, J. et al. Effectiveness of Covid-19 Vaccines against the B.1.617.2 (Delta) Variant. N. Engl. J. Med. 385, 585–594 (2021).

8. Bruxvoort, K. J. et al. Effectiveness of mRNA-1273 against delta, mu, and other emerging variants of SARS-CoV-2: test negative case-control study. BMJ 375, e068848 (2021).

9. Premkumar, L. et al. The receptor binding domain of the viral spike protein is an immunodominant and highly specific target of antibodies in SARS-CoV-2 patients. Sci. Immunol. 5, eabc8413 (2020).

10. Wilhelm, A. et al. Reduced Neutralization of SARS-CoV-2 Omicron Variant by Vaccine Sera and Monoclonal Antibodies. MedRxiv (2021) doi:10.1101/2021.12.07.21267432.

11. Planas, D. et al. Considerable escape of SARS-CoV-2 Omicron to antibody neutralization. Nature 1–7 (2021) doi:10.1038/s41586-021-04389-z.

12. VanBlargan, L. A. et al. An infectious SARS-CoV-2 B.1.1.529 Omicron virus escapes neutralization by therapeutic monoclonal antibodies. Nat. Med. 1–6 (2022) doi:10.1038/s41591-021-01678-y.

13. Hoffmann, M. et al. The Omicron variant is highly resistant against antibody-mediated neutralization: Implications for control of the COVID-19 pandemic. Cell 185, 447-456.e11 (2022).

14. Garcia-Beltran, W. F. et al. mRNA-based COVID-19 vaccine boosters induce neutralizing immunity against SARS-CoV-2 Omicron variant. Cell S0092-8674(21)01496–3 (2022) doi:10.1016/j.cell.2021.12.033.

15. Sheward, D. J. et al. Variable loss of antibody potency against SARS-CoV-2 B.1.1.529 (Omicron). BioRxiv (2021) doi:10.1101/2021.12.19.473354.

16. Gray, G. E. et al. Vaccine effectiveness against hospital admission in South African health care workers who received a homologous booster of Ad26.COV2 during an Omicron COVID19 wave: Preliminary Results of the Sisonke 2 Study. MedRxiv (2021) doi:10.1101/2021.12.28.21268436.

17. GeurtsvanKessel, C. H. et al. Divergent SARS CoV-2 Omicron-reactive T- and B cell responses in COVID-19 vaccine recipients. Sci. Immunol. (2022) doi:10.1126/sciimmunol.abo2202.

18. Liu, J. et al. Vaccines Elicit Highly Conserved Cellular Immunity to SARS-CoV-2 Omicron. Nature 1–7 (2022) doi:10.1038/s41586-022-04465-y.

19. Keeton, R. et al. T cell responses to SARS-CoV-2 spike cross-recognize Omicron. Nature 1–5 (2022) doi:10.1038/s41586-022-04460-3.

20. Richardson, S. I. et al. SARS-CoV-2 Omicron triggers cross-reactive neutralization and Fc effector functions in previously vaccinated, but not unvaccinated individuals. MedRxiv (2022) doi:10.1101/2022.02.10.22270789.

21. Jennewein, M. F. et al. Isolation and Characterization of Cross-Neutralizing Coronavirus Antibodies from COVID-19+ Subjects. BioRxiv (2021) doi:10.1101/2021.03.23.436684.

22. Liu, P. et al. Novel ACE2-Fc chimeric fusion provides long-lasting hypertension control and organ protection in mouse models of systemic renin angiotensin system activation. Kidney Int. 94, 114–125 (2018).

23. Solforosi, L. et al. Immunogenicity and efficacy of one and two doses of Ad26.COV2.S COVID vaccine in adult and aged NHP. J. Exp. Med. 218, e20202756 (2021).

24. Jongeneelen, M. et al. Ad26.COV2.S elicited neutralizing activity against Delta and other SARS-CoV-2 variants of concern. BioRxiv (2021) doi:10.1101/2021.07.01.450707.

25. Han, P. et al. Receptor binding and complex structures of human ACE2 to spike RBD from omicron and delta SARS-CoV-2. Cell 185, 630-640.e10 (2022).

26. Mannar, D. et al. SARS-CoV-2 Omicron variant: Antibody evasion and cryo-EM structure of spike protein–ACE2 complex. Science 375, 760–764 (2022).

27. Collie, S., Champion, J., Moultrie, H., Bekker, L.-G. & Gray, G. Effectiveness of BNT162b2 Vaccine against Omicron Variant in South Africa. N. Engl. J. Med. 386, 494–496 (2022).

28. UK Health Security Agency. SARS-CoV-2 variants of concern and variants under investigation in England Technical briefing: Update on hospitalisation and vaccine effectiveness for Omicron VOC-21NOV-01 (B.1.1.529). https://assets.publishing.service.gov.uk/government/uploads/system/uploads/attachment_data/file/1045619/Technical-Briefing-31-Dec-2021-Omicron_severity_update.pdf (2021).

29. Chandrashekar, A. et al. Vaccine Protection Against the SARS-CoV-2 Omicron Variant in Macaques. BioRxiv (2022) doi:10.1101/2022.02.06.479285.

30. Lee, I.-J. et al. Omicron-specific mRNA vaccine induced potent neutralizing antibody against Omicron but not other SARS-CoV-2 variants. (2022) doi:10.1101/2022.01.31.478406.

31. Ying, B. et al. Boosting with Omicron-matched or historical mRNA vaccines increases neutralizing antibody responses and protection against B.1.1.529 infection in mice. BioRxiv (2022) doi:10.1101/2022.02.07.479419.

32. Wu, Y. et al. Omicron-specific mRNA vaccine elicits potent immune responses in mice, hamsters, and nonhuman primates. BioRxiv (2022) doi:10.1101/2022.03.01.481391.

33. Zang, J. et al. An mRNA vaccine candidate for the SARS-CoV-2 Omicron variant. BioRxiv (2022) doi:10.1101/2022.02.07.479348.

34. Hawman, D. W. et al. Replicating RNA platform enables rapid response to the SARS-CoV-2 Omicron variant and elicits enhanced protection in naïve hamsters compared to ancestral vaccine. BioRxiv (2022) doi:10.1101/2022.01.31.478520.

35. Gagne, M. et al. mRNA-1273 or mRNA-Omicron boost in vaccinated macaques elicits comparable B cell expansion, neutralizing antibodies and protection against Omicron. (2022) doi:10.1101/2022.02.03.479037.

36. He, X. et al. A homologous or variant booster vaccine after Ad26.COV2.S immunization enhances SARS-CoV-2-specific immune responses in rhesus macaques. Sci. Transl. Med. eabm4996 (2022) doi:10.1126/scitranslmed.abm4996.

37. Yamasoba, D. et al. Virological characteristics of SARS-CoV-2 BA.2 variant. BioRxiv (2022) doi:10.1101/2022.02.14.480335.

